# The evolution of interdependence in a four-way mealybug symbiosis

**DOI:** 10.1101/2021.01.28.428658

**Authors:** Arkadiy I. Garber, Maria Kupper, Dominik R. Laetsch, Stephanie R. Weldon, Mark S. Ladinsky, Pamela J. Bjorkman, John P. McCutcheon

## Abstract

Mealybugs are insects that maintain intracellular bacterial symbionts to supplement their nutrient-poor plant sap diets. Some mealybugs have a single betaproteobacterial endosymbiont, a *Candidatus* Tremblaya species (hereafter *Tremblaya*) that alone provides the insect with its required nutrients. Other mealybugs have two nutritional endosymbionts that together provide these nutrients, where *Tremblaya* has gained a gammaproteobacterial partner that resides in the cytoplasm of *Tremblaya*. Previous work had established that *Pseudococcus longispinus* mealybugs maintain not one but two species of gammaproteobacterial endosymbionts along with *Tremblaya*. Preliminary genomic analyses suggested that these two gammaproteobacterial endosymbionts have large genomes with features consistent with a relatively recent origin as insect endosymbionts, but the patterns of genomic complementarity between members of the symbiosis and their relative cellular locations were unknown. Here, using long-read sequencing and various types of microscopy, we show that the two gammaproteobacterial symbionts of *P. longispinus* are mixed together within *Tremblaya* cells, and that their genomes are somewhat reduced in size compared to their closest non-endosymbiotic relatives. Both gammaproteobacterial genomes contain thousands of pseudogenes, consistent with a relatively recent shift from a free-living to endosymbiotic lifestyle. Biosynthetic pathways of key metabolites are partitioned in complex interdependent patterns among the two gammaproteobacterial genomes, the *Tremblaya* genome, and horizontally acquired bacterial genes that are encoded on the mealybug nuclear genome. Although these two gammaproteobacterial endosymbionts have been acquired recently in evolutionary time, they have already evolved co-dependencies with each other, *Tremblaya*, and their insect host.

**Significance:** Mealybugs are sap-feeding insects that house between one and three bacterial endosymbionts to supplement their nutritionally poor diets. Many mealybug-bacteria relationships were established tens or hundreds of millions of years ago, and these ancient examples show high levels host-endosymbiont genomic and metabolic integration. Here, we describe the complete genomes and cellular locations for two bacterial endosymbiont which have recently transitioned from a free-living to an intracellular state. Our work reveals the rapid emergence of metabolic interdependence between these two nascent endosymbionts, their partner bacterial co-symbiont in whose cytoplasm they reside, and their insect host cell. Our work confirms that intracellular bacteria rapidly adapt to a host-restricted lifestyle through breakage or loss of redundant genes.

## Introduction

Insects with nutrient-poor diets (e.g. plant sap, blood, wood) retain nutritional symbionts that supplement their diet with nutrients, such as amino acids and vitamins (Baumann, 2005; Douglas, 2006). Mealybugs (**Figure 1A**) are insects that exclusively consume phloem sap and maintain nutritional endosymbiotic bacteria within specialized cells called bacteriocytes (Buchner, 1965; von Dohlen *et al*., 2001; Baumann *et al*., 2002). Mealybug bacteriocytes house between one and three different bacterial endosymbionts depending on the mealybug species (Kono *et al*., 2008; Koga *et al*., 2013; López-Madrigal *et al*., 2013; Husník and McCutcheon, 2016; Szabó *et al*., 2017; Gil *et al*., 2017). These mealybug endosymbionts produce essential amino acids and vitamins, which are present at low and variable levels in the insect’s specialized plant sap diet. While it is not uncommon for insects to simultaneously maintain multiple endosymbionts (Buchner, 1965; Fukatsu *et al*., 1998; Toh *et al*., 2006; Thao *et al*., 2002; Moran *et al*., 2008; McCutcheon and Moran, 2010), the spatial organization of the dual mealybug endosymbionts is unusual: each bacteriocyte houses cells of *Candidatus* Tremblaya princeps (Betaproteobacteria; hereafter referred to as *Tremblaya*), and inside each *Tremblaya* cell reside tens to hundreds of cells of another endosymbiont from the gammaproteobacterial family Enterobacteriaceae (von Dohlen *et al*., 2001). Many of these intra-Tremblaya endosymbionts are members of the *Sodalis* genus, which are commonly found as endosymbionts of insects (Oakeson *et al*., 2014; Clayton *et al*., 2012; Hall *et al*., 2020; Toh *et al*., 2006 Husník and McCutcheon, 2016; McCutcheon *et al*., 2019).

**Figure 1:**
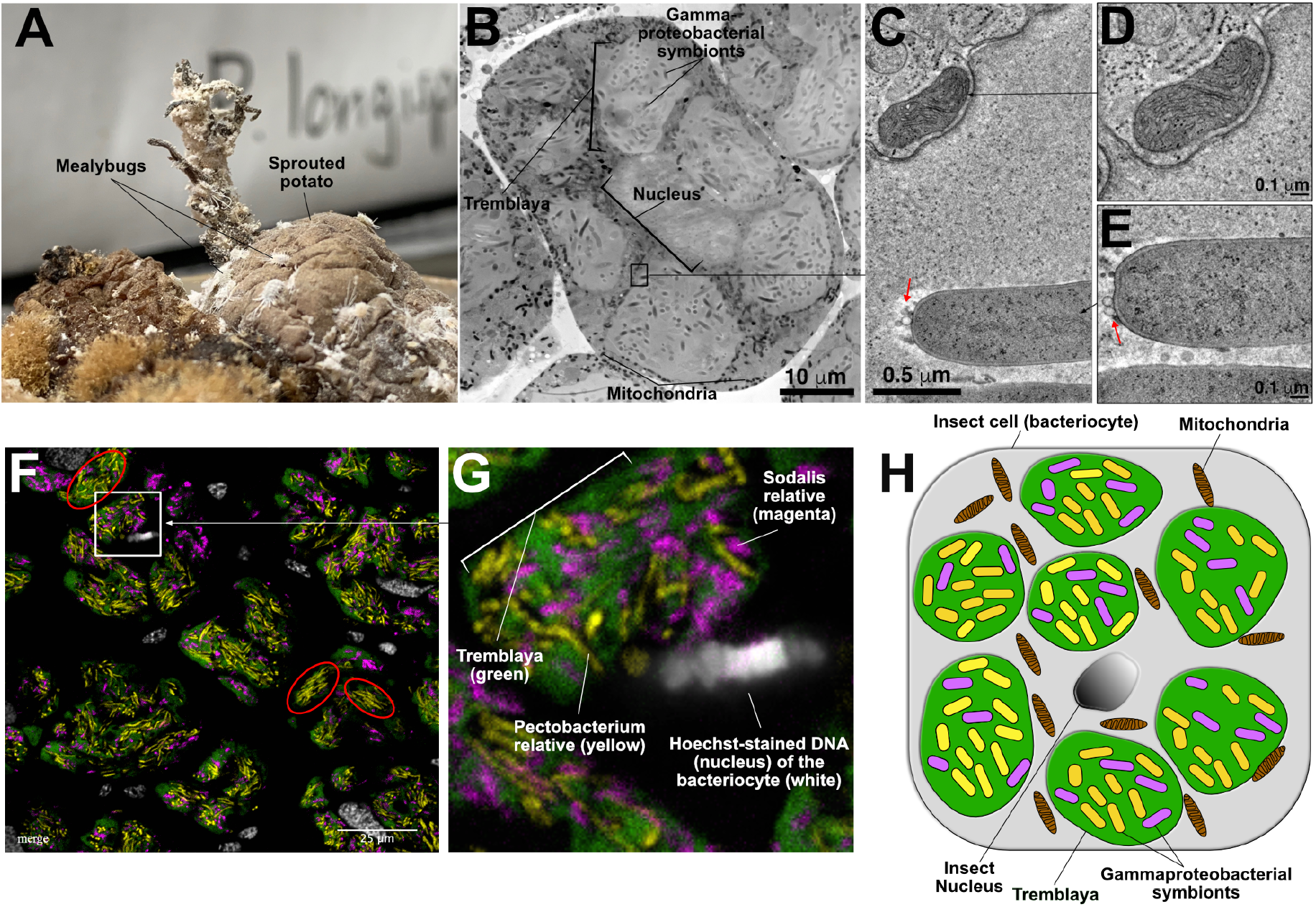
The structure of the P. longispinus symbiosis. **A)** Image of P. longispinus mealybugs on a sprouted potato. **B)** Montaged TEM overview image of a bacteriocyte from P. longispinus. The 6-7 light gray blobs are Tremblaya cells, surrounding a central eukaryotic nucleus. Within each Tremblaya cell reside rod-shaped and more electron-dense gammaproteobacterial cells. Black-colored rods in between Tremblaya are mitochondria within eukaryotic cytoplasm. The insect nucleus is at the center of the bacteriocyte in a gray shade that’s similar to Tremblaya. **C)** Detail from an electron tomographic slice showing the boundary of a Tremblaya cell, where a mitochondrion is visible near the Tremblaya cell envelope. **D)** Higher magnification view of the mitochondrion shown in B. **E)** Tomographic slice of a gammaproteobacterial symbiont that resides inside Tremblaya, showing numerous outer membrane vesicles (red arrows). The bacterial symbionts are easily distinguished from eukaryotic mitochondria. **F)** Fluorescent in situ hybridization (FISH) image of P. longispinus bacteriome tissue showing the localization of two different gammaproteobacterial endosymbionts within Tremblaya cells. Fluorophore-labelled probes were used to localize Tremblaya cells (green) and the two gammaproteobacterial endosymbionts (yellow and magenta). DNA and therefore insect nuclei were counterstained with Hoechst (white). They appear to each be surrounded by several Tremblaya cells per bacteriocyte. **G)** Zoomed in and annotated detail fluorescence microscopy image of a P. longispinus bacteriocyte. **H)** Schematic representation of P. longispinus bacteriocytes.

Genomic studies of numerous insect-endosymbiont systems have revealed strong and consistent patterns of complementary gene loss and retention among all members of the symbiosis (Shigenobu *et al*., 2000; van Ham *et al*., 2003; Wu *et al*., 2006; Gatehouse *et al*., 2012; Sloan and Moran, 2012; McCutcheon and Moran, 2010; Łukasik *et al*., 2018). While in most cases, a single endosymbiont genome will retain complete or near-complete pathways for individual metabolites, mealybug endosymbionts are unusual in that the reciprocal pattern of gene loss and retention exists *within* biochemical pathways (McCutcheon and von Dohlen, 2011; Husník 2013; López-Madrigal *et al*., 2013; Husník 2016; Szabó *et al*., 2017; Gil *et al*., 2017). Most of the previously published mealybug endosymbiont genomes were highly reduced in size (less than 1 Mb) and gene dense (containing few pseudogenes), which made discerning these complementary gene loss and retention patterns relatively straightforward.

*P. longispinus* harbors the symbiont *Tremblaya*, but unlike *Tremblaya* in other mealybugs, the *P. longispinus* strain of *Tremblaya* houses not one but two species of gammaproteobacterial endosymbionts (Gatehouse *et al*., 2011; Rosenblueth *et al*., 2012; Husník and McCutcheon, 2016). We previously reported draft genome assemblies of these two gammaproteobacterial endosymbionts, which suggested that their combined genome sizes were large, approximately 8.2 megabase pairs (Mbp) in length (Husník and McCutcheon, 2016). Phylogenetic analysis showed that one of these gammaproteobacterial symbionts belonged to the *Sodalis* genus, and the other was more closely related to members of the *Pectobacterium* genus. However, the poor quality of these draft genome assemblies made detailed genomic analysis impossible. Light microscopy on *P. longispinus* (Gatehouse *et al*., 2012) suggested that the gammaproteobacterial endosymbionts resided inside *Tremblaya* cells, as is the case in other mealybugs (von Dohlen *et al*., 2001). But it was unclear from these data whether 1) one or both of these gammaproteobacteria were restricted to *Tremblaya* cells (that is, if they were also found in the cytoplasm of the host insect bacteriocyte), 2) whether each gammaproteobacterial species was restricted to particular *Tremblaya* cell types, or 3) whether the two gammaproteobacterial symbionts were mixed together inside undifferentiated *Tremblaya* cells. Here, we add long-read data generated from *P. longispinus* bacteriome tissue to greatly improve the gammaproteobacterial genome assemblies and annotations. We describe the relative cellular locations of the endosymbionts using fluorescence and transmission electron microscopy, and report the genome evolutionary patterns and metabolic contributions of the microbial members of this unusual four-way symbiosis.

## Material and Methods

### Insect rearing

*Pseudococcus longispinus* populations were reared on sprouted potatoes (*Figure 1A*) at 25C, 77% relative humidity, and a 12h light/dark cycle in a Percival 136LL incubator.

### *RNA Fluorescence* in situ *Hybridization (RNA-FISH)*

Whole *P. longispinus* individuals of the second and third instar developmental stage were submerged in Ringer solution (3 mM CaCl2 * 2H2O, 182 mM KCl, 46 mM NaCl, 10 mM Tris base; adjusted to pH 7.2) and carefully opened for better buffer infiltration. Samples were transferred into Carnoy’s fixative (EtOH: chloroform: acetic acid; 6:3:1) and fixed overnight at 4°C. Tissue samples were then dehydrated in a graded ethanol series from 70% to 100% ethanol. Samples were transferred into tissue bags and cassettes for paraffin embedding using a Leica ASP 300 Tissue Processor. Ethanol was exchanged for methyl salicylate, and then incubated in 100% xylene before infiltration with paraffin. Each individual sample was embedded in a single paraffin block, semithin sections (5-6μm) were prepared with a microtome and mounted onto microscopy slides.

Sections designated for RNA-FISH experiments were deparaffinized in xylene and rehydrated in a graded ethanol series (100% to 30%). Tissue sections were then prehybridized in hybridization buffer (900mM NaCl, 20mM Tris-HCl pH 7.5, 35% formamide). Hybridization was performed by adding 1.5 – 2 μL of the probe targeting the *Pectobacterium*-related endosymbiont (5’[Cy3]-ccacgcctcaagggcacaacctc; 100 μM) to each 100 μL hybridization buffer and incubated at 40C. Samples were then briefly rinsed in wash buffer (70 mM NaCl, 20 mM Tris-HCl pH 7.5, 5 mM EDTA pH 8.0, 0.01% SDS) before mounting the slides with hybridization buffer supplemented with 1.5 – 2 μL of each probe targeting the *Sodalis*-related endosymbiont (5’[Cy5]-aaagccacggctcaaggccacaacctt; 100 μM) and *Tremblaya* (5’[fluorescein]-gccttagcccgtgctgccgtac; 100 μM) per 100μl buffer, followed by overnight incubation at 30C. Slides were then washed in washing buffer at 30°C and counterstained with Hoechst in washing buffer. After another washing step, sample slides were rinsed in dH2O, mounted with FluorSave™ Reagent (Sigma Millipore), and analyzed by confocal laser scanning microscopy with a Zeiss LSM 880. Images were processed using Fiji version 1.0.

### Electron Microscopy

Bacteriomes were dissected from P. longispinus individuals as previously described (Bublitz *et al*., 2019). Isolated bacteriomes were pre-fixed with 3% glutaraldehyde, 1% paraformaldehyde, 5% sucrose in 0.1M sodium cacodylate trihydrate for 12 – 24 h at 4°C, then rinsed briefly with cacodylate buffer. Bacteriomes were placed into brass planchettes (Ted Pella, Inc.) pre-filled with cacodylate buffer + 10% 70kD Ficoll (extracellular cryoprotectant; Sigma) and ultra-rapidly frozen with a HPM010 High Pressure Freezing machine (Bal-Tec/ABRA, Switzerland). Vitreously frozen samples were transferred under liquid nitrogen to Nunc cryovials (Thermo-Fisher Scientific) containing 2% OsO_4_, 0.05% uranyl acetate in acetone and placed into an AFS-2 Freeze Substitution Machine (Leica Microsystems, Austria). Samples were freeze-substituted at −90°C for 72 h, warmed to −20°C over 12 h, held at −20°C for an additional 12 h and then brought to room temperature. Samples were rinsed 4x with acetone, infiltrated into Epon-Araldite resin (Electron Microscopy Sciences, Port Washington PA), then flat-embedded between two Teflon-coated glass microscope slides. Resin was polymerized at 60°C for 24-48 h. Embedded samples were observed by phase-contrast microscopy to ascertain specimen quality and to select appropriate regions for EM study. Blocks of tissue (typically containing a single bacteriome) were excised with a scalpel and glued to plastic sectioning stubs. Serial semi-thick (150 – 300 nm) sections were cut with a UC6 ultramicrotome (Leica Microsystems) using a diamond knife (Diatome Ltd, Switzerland) and collected onto Formvar-coated copper-rhodium 1 mm slot grids (Electron Microscopy Sciences). Grids were stained with 3% uranyl acetate and lead citrate, then 10 nm colloidal gold particles were applied to both sides of the sections to serve as fiducial markers for subsequent tomographic image alignment.

### Dual Axis Tomography

Grids were placed in a Dual-Axis tomography specimen holder (Model 2040; E.A. Fischione Instruments Inc., Export PA) and viewed with a Tecnai TF-30ST transmission electron microscope at 300K eV. Dual-axis tilt-series were acquired automatically using the SerialEM software package (Mastronarde., 2005) and recorded digitally with a 2k x 2k CCD camera (XP1000; Gatan, Inc. Pleasanton CA). Briefly, sections were tilted +/- 64° with images taken at 1° increments. The grid was then rotated 90° and a similar tilt-series was recorded around the orthogonal axis. Tomograms were calculated, joined and analyzed using the IMOD software package (Mastronarde., 2008; Mastronarde and Held, 2017) on MacPro and iMac Pro computers (Apple Inc.).

### Sequencing and Assembly

Raw Illumina HiSeq 2000 reads published in (Husník and McCutcheon, 2106; BioProject: PRJEB12068) were downloaded from the National Center for Biotechnology Information (NCBI) Sequence Read Archive (SRA), using the SRA Toolkit v2.10.8 (SRA Toolkit Development Team). Reads were trimmed using Trimmomatic v0.36 (minimum length=36 bp, sliding window=4 bp, minimum quality score=15 [ILLUMINACLIP:TruSeq3-PE:2:30:10 LEADING:3 TRAILING:3 SLIDINGWINDOW:4:15 MINLEN:36]) (Bolger *et al*., 2014). For PacBio sequencing, genomic DNA was prepared from pooled mealybugs using Qiagen Genomic Tip 500 g extraction kits, size selected for fragments >20 kb using a BluePippen device, followed by library preparation using a SMRTbell Template Prep Kit v1.0. The resulting libraries were sequenced on 28 single molecule realtime (SMRT) PacBio cells using P6 version 2 chemistry and reagents by Sci-Life labs in Uppsala, Sweden. This sequencing effort resulted in 6,101,355 reads of average length 9,805 bases for a total of 59,828,022,374 bases. These reads were error corrected and trimmed, resulting in 5.05 million reads with average sequence length of 9,318 bases. These corrected and trimmed reads were then assembled in Canu v1.6 (correctedErrorRate=0.45, genomesize=284m) (Koren *et al*., 2017), which produced 3,049 contigs spanning 438,113,873 bases. Preliminary gammaproteobacterial contigs were extracted from the Canu assembly using the SprayN-Pray software (https://github.com/Arkadiy-Garber/SprayNPray). Briefly, SprayNPray predicts open reading frames (ORFs) using Prodigal (Hyatt *et al*., 2010), and then queries each ORF against NCBI’s RefSeq database (release 200) (Pruitt *et al*., 2007) using DIAMOND v2.0.4.142 (Buchfink *et al*., 2014, e-value 1E-6). Putative endosymbiont contigs were then extracted from the larger assembly based on gene density, GC-content, and taxonomy of top DIAMOND hits (to *Sodalis*- and *Pectobacterium/Brenneria*-related spp.) to each contig. These contigs were then used to identify and extract all Illumina and PacBio reads associated with the gammaproteobacterial symbionts. Identification of endosymbiont-affiliated Illumina reads was performed using Bowtie2 v2.3.4.1 (Langmead and Salzberg, 2012). Identification of endosymbiont PacBio reads was performed using BLASR v5.1 (Chaisson and Tesler, 2012). About 3.2% of all PacBio reads mapped to the CANU-assembled contigs affiliated with the gammaproteobacterial endosymbionts. Of the 124.5 million Illumina read pairs, 4.8% mapped to the crude gammaproteobacterial endosymbiont contigs.

Once these gammaproteobacterial subsets of short and long reads were identified and extracted, Unicycler v0.4.8 (Wick *et al*., 2017) was used, with “normal” mode (minimum bridge quality = 10) to carry out a hybrid SPAdes v3.13.0 (Bankevich *et al*., 2012) assembly (default k-mers). The two gammaproteobacterial genomes were binned using a combination of metrics, including coverage of Illumina reads mapped against the final assemblies. Coverage of Bowtie2-mapped Illumina reads was estimated using the jgi_summarize_bam_contig_depths script from the MetaBAT package (Kang *et al*., 2019). Since the closest phylogenomic affiliations of each gammaproteobacterial symbiont are known (Husník and McCutcheon, 2016), we also used BLASTP (Camacho *et al*., 2009) to compare the open reading frames (ORFs) from each contig against NCBI’s RefSeq database (Pruitt *et al*., 2007). ORFs from each contig were predicted using Prodigal v2.6.3 (Hyatt *et al*., 2010), and the phylogenetic affiliation of each contigs’ ORFs was inferred by its top BLASTP hit from NCBI’s RefSeq database (Pruitt *et al*., 2007).

### Phylogenomic analysis

Phylogenomic analysis was carried out using GToTree v1.5.38 (Lee *et al*., 2019) and RAxML (Stamatakis, 2014). Briefly, single-copy genes were identified using a set of HMMs for genes common to Gammaproteobacteria (Lee, 2019). As part of the GToTree pipeline, single-copy genes are identified using HMMER v3.2.1 (Johnson *et al*., 2010), aligned with Muscle v3.8 (Edgar, 2004), and concatenated. These concatenated alignments were then used to build a phylogenomic tree with RAxML, with 100 bootstraps (-N 100), the PROTCAT model for amino acid substitution, and the BLOSUM 62 amino acid matrix (-m PROTCATBLOSUM62) (Stamatakis, 2014).

### Annotation and Biosynthetic Pathway Reconstruction

Each endosymbiont was annotated using Prokka v1.14.6 (Seemann, 2014). As part of Prokka’s pipeline, coding regions are detected using Prodigal; noncoding RNA sequences were also identified: tRNAs and tmRNAs using Aragorn (Laslett and Canbeck, 2004), and rRNAs using RNAmmer (Lageson *et al*., 2007). Prokka annotation also included identification of transposases, using the ISfinder database of insertion sequences (Siguier *et al*., 2006). Genes were also annotated using the GhostKOALA v.2.2 web server, which uses the Kyoto Encyclopedia of Genes and Genomes (KEGG) Orthology database (Kenehisa *et al*., 2016). Biosynthetic pathways for amino acids, vitamins, peptidoglycan, and translation-related genes were manually identified from these annotations and organized into pathways. Horizontal gene transfers (HGTs) present on the mealybug genome were previously identified (Husník and McCutcheon, 2016; Bublitz, *et al*., 2019).

### Pseudogene Prediction

Candidate pseudogenes were identified with the Pseudofinder software (https://github.com/filip-husnik/pseudofinder), using DIAMOND (–diamond) (Buchfink *et al*., 2014) to find each ORF’s closest homologs NCBI’s RefSeq (Pruitt *et al*., 2007) database. This allowed us to identify pseudogenes based on length and gene fragmentation due to early stop codons. ORFs that deviate more than 25% (–length_pseudo 75) from the average length of the 15 top homologs (–hitcap 15, –evalue 1E-4) from RefSeq were flagged as potential pseudogenes. Additionally, ORFs with stop codons and frame-shift mutations were also flagged as pseudogenes. These fragmented ORFs were identified by Pseudofinder by finding adjacent ORFs that have the same gene as their top DIAMOND hit. For adjacent ORFs to be considered as fragmented parts of the same ancestral gene, we used a distance cutoff of 2000 bp (–distance 2000). The length of each gene and pairwise homology was also taken into consideration to exclude intact adjacent ORFs that represent gene duplication events resulting in tandem-encoded duplicate ORFs. Non-genic regions, in which Prodigal did not detect any ORFs were also compared against NCBI’s Ref-Seq database, to identify pseudogenized ORFs that no longer appear as ORFs to Prodigal’s algorithm. Non-genic regions required at least five DIAMOND matches to proteins in the Ref-Seq database (–intergenic_threshold 0.3) to be considered pseudogenes.

Using DIAMOND BLASTP, Pseudofinder also compared ORFs from each endosymbiont to its closest ancestor, inferred from phylogenomic analysis and average amino acid identity. We identified *Pectobacterium wasabiae* as the closest free-living relative for one endosymbiont and *Sodalis praecaptivus* HS for the other. Using PAL2NAL v14 (Suyama *et al*., 2006), Pseudofinder generates codon alignments for each ortholog pair, then, using Codeml v4.9j (Yang *et al*., 2007), calculates *dN/dS* values for each pairwise comparison. We provide the control file (codeml.ctl) containing the parameters used by Codeml in the following GitHub repository: https://github.com/Arkadiy-Garber/PLON-genome-paper. We required *dS* to be greater than 0.001 and lower than 3 for *dN/dS* calculation (-m 0.001, -M 3). This allowed us to infer cryptic pseudogenes, or genes that are likely undergoing relaxed selection but have not acquired any obvious inactivating mutations (Clayton *et al*., 2012; Oakeson *et al*., 2014; Van Leuven *et al*., 2014). We used a *dN/dS* cutoff of 0.3 (Oakeson *et al*., 2014), flagging genes as pseudogenes if their *dN/dS* values are higher than this threshold (–max_dnds 0.3).

Pseudogene calls from Pseudofinder were manually inspected, using AliView (Larsson, 2014) to confirm gene fragmentation, *dN/dS* values, and other inactivating mutations.

### Identification of duplicated genes

We used ParaHunter to identify gene duplicates in the endosymbiont genomes (Miller *et al*, 2020). This software uses MMseqs2 v12.113e3 (Steinegger and Söding, 2017) to identify homologous gene clusters within each genome. For this analysis, we used a cutoff of 50% amino acid identify (-m 0.5) over at least 50% of the length (-l 0.5) of target sequence. After clusters are identified, within-cluster analysis is carried out, where pairwise amino acid and nucleotide alignments are converted to codon alignments using PAL2NAL (Suyama *et al*., 2006), and then *dN/dS* is calculated using Codeml (Yang *et al*., 2007). Estimation of *dN/dS* required *dS* values greater than 0.001 and lower than 3.

### Additional scripts and plotting

Additional custom python scripts were used to process the data presented in this study. These scripts are all annotated and available in the following GitHub repository: https://github.com/Arkadiy-Garber/PLON-genome-paper. Many plots presented in this study were made in R (R Core Team, 2013), using the following packages: ggplot2 (Wickham, 2009) and reshape (Wickham, 2007).

### Data Availability

PacBio reads were deposited to the Sequence Read Archive (SRA), under NCBI BioProject PRJNAXXXXXX. Genome assemblies for the gammaproteobacterial endosymbionts are available under NCBI BioProject PRJNAXXXXXX. Genome sequences and annotation data for the two gammaproteobacterial endosymbionts were also made available via figshare: Genome sequence and Prokka-annotation are available at https://doi.org/10.6084/m9.figshare.13632407.v1 for the *Pectobacterium*-related symbiont and https://doi.org/10.6084/m9.figshare.13632398.v2 for the *Sodalis*-related symbiont. Pseudogene predictions are available at https://doi.org/10.6084/m9.figshare.13632419.v1 for the *Pectobacterium*-related symbiont and https://doi.org/10.6084/m9.figshare.13632416.v1 for the *Sodalis*-related symbiont. Files used in the analysis of other *Sodalis*- and *Symbiopectobacterium*-related endosymbionts are available at the following address: https://doi.org/10.6084/m9.figshare.13661189.

## Results

### Gammaproteobacterial endosymbionts are located within Tremblaya

Previous light microscopy suggested that at least some, if not all, of the gammaproteobacterial endosymbionts of *P. longispinus* resided inside of *Tremblaya* (Gatehouse *et al*., 2011). However, these data lacked the resolution to clarify whether or not both species of gammaproteobacterial endosymbionts were exclusively contained within *Tremblaya*, or whether some might also live in the cytoplasm of bacteriocytes. We used transmission electron microscopy (TEM) to identify the localization of the gammaproteobacterial cells. Our TEM data suggest that all gammaproteobacterial endosymbiont cells are contained within *Tremblaya* and are not free in the cytoplasm of the host insect cell (**Figure 1B-E**). At low magnification, elongated cells of different shapes and sizes, which we presume to be the gammaproteobacterial symbionts, can be seen inside of *Tremblaya* cells (**Figure 1B**). Structures of similar size and electron density can be observed outside of *Tremblaya* cells, but these were found to be mitochondria upon examination at higher magnification (**Figure 1C-D**). We note numerous outer-membrane vesicles (OMVs) apparently being extruded by the *Sodalis*- or *Pectobacterium*-related endosymbiont cells (**Figure 1E**) (Toyofuku *et al*., 2019). The function of these OMVs in the symbiosis, if any, is unknown.

Using the small subunit (SSU) ribosomal RNA sequences reported in (Husník and McCutcheon, 2016), we next performed fluorescence *in situ* hybridization (FISH) targeting SSU rRNA to establish the relative locations of the two gammaproteobacterial endosymbionts. Here, we were testing whether there were two different types of *Tremblaya* cells, each containing only one type of gammaproteobacterial cell, or whether both gammaproteobacterial species were mixed together inside of one type of *Tremblaya* cell. We find that both gammaproteobacterial endosymbionts are mixed together in one type of *Tremblaya* cell (**Figure 1F-H**). The overall distribution of endosymbionts within *Tremblaya* cells suggests that the *Pectobacterium* relative is more abundant than the *Sodalis* relative (colored yellow and violet, respectively, in **Figure 1F-G**). We note that there are some *Tremblaya* cells or regions of *Tremblaya* cells where the *Pectobacterium* relative appears highly abundant, with almost no cells of the *Sodalis* relative present (red circles in **Figure 1F**). Cells of the *Pectobacterium* relative appear to be longer than cells of the *Sodalis* relative. This mixture of long and short cells is consistent with what we see in the TEM images, although the identities of the two cell types cannot be discerned in TEM (**Figure 1B**).

### Gammaproteobacterial endosymbionts have large genomes similar to free-living bacteria

Previous efforts to assemble the genomes of the two gammaproteobacterial symbionts using only short read Illumina technology resulted in highly fragmented genome assemblies (Husník and McCutcheon, 2016). Our addition of PacBio reads greatly improved the quality of the *Sodalis* symbiont’s genome (3 contigs vs 200 contigs from short reads alone), likely due to their ability to span repetitive insertion sequences (IS) that appear to be abundant in the *Sodalis* endosymbiont genome (**Supplemental File 1**). The *Pectobacterium*-related symbiont genome was also improved by long-read sequencing (9 vs 40 contigs from short reads alone). To aid in the binning of contigs for each symbiont genome, we relied on the differential read coverage calculated from Illumina short reads for the two symbiont genomes as well as the similarity of genes compared to non-endosymbiont *Sodalis* and *Pectobacterium* genomes. Despite numerous computational and PCR-based experiments, we were unable to close either genome into complete circular-mapping molecules.

At 4.3 Mb and 3.6 Mb, both gammaproteobacterial endosymbiont genomes are similar in size to genomes of many free-living bacteria. One circular-mapping contig of 150 kb was putatively assigned as a plasmid to the *Pectobacterium* relative, based on its circular structure and its ORFs showing high similarity to genes from other *Pectobacterium* and *Brenneria* spp. One additional contig of 90 kb containing genes with high similarity to other *Sodalis* spp. genes showed 3-fold higher coverage relative to the other *Sodalis*-related contigs; this could either be a plasmid, or a large repeat region of the genome. Finally, we identified a plasmid seemingly related to other *Arsenophonus* plasmids, but we were unable to associate it to either gammaproteobacterial endosymbiont. An overview of the endosymbiont genomes is shown in **Table 1**.

**Table 1.** Assembly summary for the three endosymbionts and three putative plasmids assembled from *P. longispinus* bacteriomes. The *Pectobacterium* and *Sodalis* symbionts are named *Symbiopectobacterium endolongispinus* and *Sodalis endolongispinus*, respectively, as discussed in the section “ ***Naming of the two gammaproteobacterial symbionts***” *Genome of *Tremblaya* taken from Husník and McCutcheon, 2016.

| Endosymbionts and plasmids | Genome/plasmid size (bp) | Average read depth (Illumina) | Number of contigs |
| --- | --- | --- | --- |
| *Pectobacterium* symbiont | 4,343,494 | 43.08 | 9 |
| *Pectobacterium* plasmid | 148,954 | 47.04 | 1 |
| *Sodalis* symbiont | 3,638,256 | 22.33 | 3 |
| *Sodalis* (possible plasmid) | 89,872 | 56.70 | 1 |
| *Tremblaya* * | 144,042 | 1,565.37 | 1 |
| *Arsenophonus* plasmid (unbinned) | 64,583 | 160.98 | 1 |

Mapping of Illumina sequence reads to the endosymbiont genomes indicates that the *Pectobacterium* endosymbiont is twice as abundant as the *Sodalis* endosymbiont. This is consistent with FISH images where the *Pectobacterium*-related cells appear more abundant than the *Sodalis*-related cells (**Figure 1B**). Illumina read mapping to the genome of *Tremblaya* suggests that it is 25 times more abundant than its gammaproteobacterial cosymbionts. It is likely that *Tremblaya’s* genome is present in hundreds or thousands of copies per cell, consistent with previous reports of extreme polyploidy in ancient endosymbionts with tiny genomes, such as *Candidatus* Hodgkinia cicadicola (Van Leuven *et al*., 2014), *Candidatus* Sulcia muelleri (Woyke *et al*., 2010), and *Buchnera aphidicola* (Komaki *et al*., 1999).

### Gammaproteobacterial symbionts are related to opportunistic pathogens known to infect insects

The closest sequenced non-endosymbiont relatives of the *P. longispinus* gammaproteobacterial symbionts are *Pectobacterium wasabiae* (average amino acid identity = 76.1%) and *Sodalis praecaptivus* HS (average amino acid identity = 86.0%; hereafter, *Sodalis* HS) (**Figure 2**). *Sodalis* HS was isolated from a human infection (Clayton *et al*., 2012; Chari *et al*., 2015), and its genome suggests that this bacterium may be an opportunistic pathogen capable of infecting animal and plant cells (Clayton *et al*., 2012). *P. wasabiae* is a known pathogen of plants and has been identified as the causative agent of potato soft rot (Gardan *et al*., 2003; Yuan *et al*., 2014; Pasanen *et al*., 2013). Phylogenomic analysis, using a concatenated set of 172 single-copy genes common to Gammaproteobacteria (Lee *et al*., 2019), confirms the affiliation of one endosymbiont squarely within the *Sodalis* genus, closely related to other recently established endosymbionts such as *Ca*. S. glossinidius (hereafter, *S. glossinidius*) and *S. pierantonius* str. SOPE (hereafter, SOPE) (Husník and McCutcheon, 2016) (**Figure 2**). Of note, SOPE was estimated to have been established as an insect endosymbiont from a *Sodalis* HS relative very recently, approximately 28,000 years ago (Clayton *et al*., 2012). Using the GC-content among 4-fold degenerate sites in the *Sodalis endosymbiont* of *P. longispinus* (in comparison to SOPE), and assuming a clock-like reduction in GC-content following host restriction, we estimate that its divergence from *Sodalis* HS occurred roughly 67,000 years ago, although we stress that this is a very rough estimate (**Supplemental File 2**).

**Figure 2:**
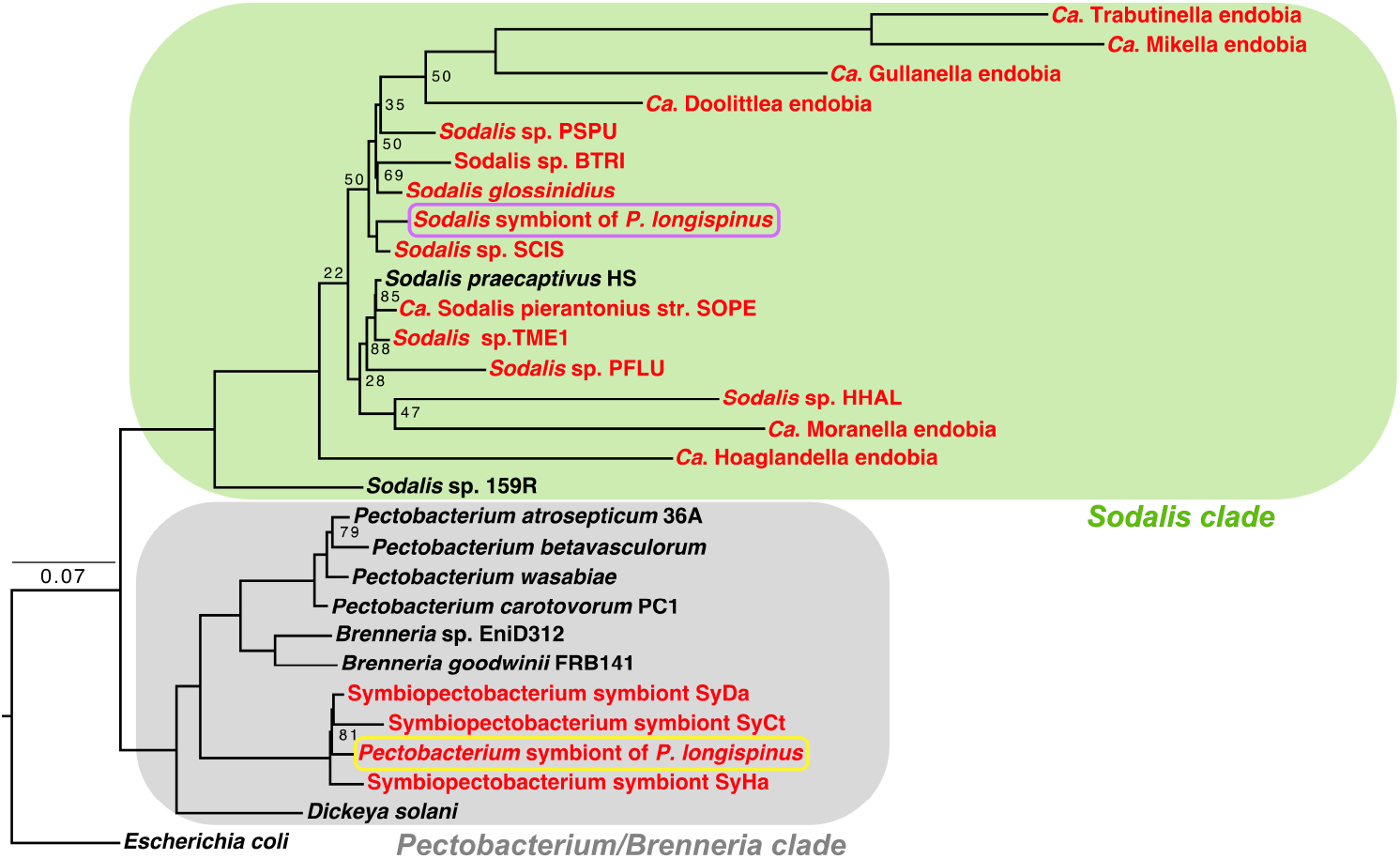
The gammaproteobacterial endosymbionts are from two different groups. A phylogenomic tree constructed with a concatenated set of 172 single-copy genes designed for Gammaproteobacteria (Lee, 2019), of Sodalis- and Pectobacterium-related endosymbionts (colored red) and the closest non-endosymbiotic relatives (colored black). Escherichia coli genome is used as the outgroup. This tree reveals two distinct clades: one containing the Pectobacterium/Brenneria-related bacteria and one containing the Sodalis-related bacteria. The two endosymbionts residing within P. longispinus bacteriocytes are emphasized in yellow (Pectobacterium-related) and violet (Sodalis-related) boxes. Unlabeled nodes have bootstrap support values greater than 90%.

The second gammaproteobacterial endosymbiont in *P. longispinus* is more closely affiliated to *Pectobacterium* and *Brenneria* spp., and appears to fall within a newly proposed group of nematode and insect endosymbionts named *Symbiopectobacterium* (Martinson *et al*., 2020). BLAST-based comparison of open-reading frames confirms that these *Symbiopectobacterium*-clade symbionts are very closely related, sharing 94-97% average nucleic acid identity (ANI) across their genomes (**Supplemental Figure 1**).

### Naming of the two gammaproteobacterial endosymbionts

For the *Sodalis* relative, we propose the name *Candidatus* Sodalis endolongispinus (hereafter, *Sod. endolongispinus*). This name highlights its close phylogenetic relationship with other bacteria in the *Sodalis* genus (**Figure 2**) and its localization inside *P. longispinus* bacteriomes. We propose the name *Candidatus* Symbiopectobacterium endolongispinus (hereafter, *Sym. endolongispinus*) for the *Pectobacterium* relative, reflecting its close phylogenetic relationship with the new *Symbiopectobacterium* group (Martinson *et al*., 2020) along with its localization inside *P. longispinus* bacteriomes.

### Pseudogenes abound in the gammaproteobacterial endosymbiont genomes

Newly established endosymbionts contain unusually high numbers of pseudogenes compared to most bacterial genomes (Toh *et al*., 2006; Burke and Moran, 2011; McCutcheon and Moran, 2012; Clayton *et al*., 2012; Oakeson *et al*., 2014). Pseudogenes are thought to form as a bacterium transitions to a strict intracellular lifecycle because many previously essential genes are no longer required in the intracellular environment (Toh *et al*., 2006; Burke and Moran, 2011; McCutcheon and Moran *et al*., 2012). Additionally, rapid pseudogenization of some genes coding for immune-stimulating compounds, such as lipopolysaccharide, is likely to be adaptive for bacteria that have recently transitioned to an intracellular lifestyle (D’Souza and Kost, 2016; McCutcheon *et al*., 2019).

The genomes of both gammaproteobacterial endosymbionts of *P. longispinus* contain thousands of pseudogenes (**Figure 3**, **Table 2**, **Supplemental Files 3-4**). The coding densities of both of these genomes are approximately 50%, much lower than average for most other free-living bacteria (Ochman and Davalos, 2006). Pseudogenes in *Sod. endolongispinus* and *Sym. endolongispinus* are found in nearly all gene categories, including membrane transport, amino acid metabolism, energy generation, secretion systems, transcriptional regulation, and motility. Several regions which appear to be remnants of prophages are also largely pseudogenized. Pseudogenes have been formed in a variety of ways, and some genes show multiple signs of pseudogenization [e.g. truncations and *dN/dS* values > 0.3 (Oakeson *et al*., 2014)] (**Figure 3B**). A substantial proportion of pseudogenes were formed by nonsense mutations, resulting in early stop codons or partial gene deletions. Consequently, many predicted pseudogenes are shorter, longer, or fragmented relative to their closest, presumably functional, homologs in non-endosymbiotic bacteria (**Figure 3C**). Many putative pseudogenes or pseudogene fragments were unrecognizable to the prokaryotic gene-finding program Prodigal (Hyatt *et al*., 2010) likely due to missing start/stop codons and/or frameshifts, and were only identified by performing BLASTX (Camacho *et al*., 2009) searches of intergenic regions against NCBI’s RefSeq database. We also detected cryptic pseudogenes, or genes that are structurally intact but are likely experiencing relaxed purifying selection, inferred from *dN/dS* ratios greater than 0.3 (Oakeson *et al*., 2014). However, we find that only a small proportion of predicted genes have elevated *dN/dS* values (0.5% of genes in *Sym. endolongispinus* and 2.2% in *Sod endolongispinus*) suggesting that most genes are still experiencing strong purifying selection (**Figure 3C**).

**Figure 3:**
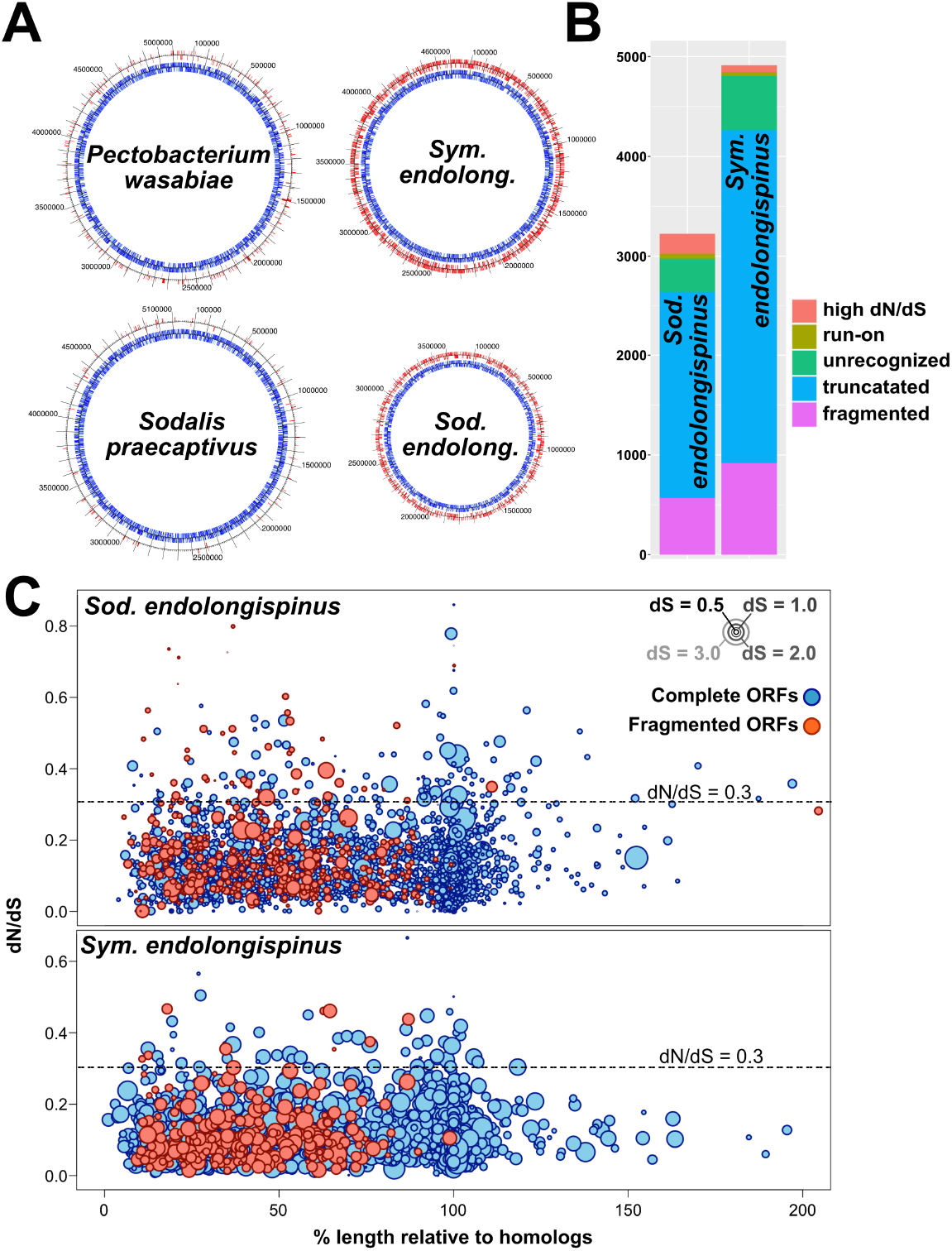
The features of gammaproteobacterial pseudogenes. **A)** Genome maps showing the positions of candidate pseudogenes in the two P. longispinus gammaproteobacterial endosymbiont genomes and their closest free-living relatives. Input genes (i.e. open reading frames [ORFs] predicted using Prokka/Prodigal) are on the inner tracks and colored blue. Predicted pseudogenes are on the outer tracks and colored red. **B)** Summary of the types of gene disruptions occurring in each of the gammaproteobacterial symbionts. The total number of disruptions is greater than the total number of pseudogenes in each genome because many pseudogenes have more than one type of disruption. **C)** Plots showing gene degradation of endosymbiont genes. Each circle represents an endosymbiont ORF; the x-axis represents the length of each gene relative to its ortholog in the reference genome (HS or P. wasabiae); the y-axis represents *dN/dS* of each gene relative to its ortholog in the reference genome; ORFs that have truncating stop codons and appear fragmented relative to orthologs in free-living genomes are colored red; finally, the size of each circle represents *dS*, a proxy for evolutionary divergence.

**Table 2.** Summary of pseudogene counts and coding densities in *Sym. endolongispinus* and *Sod. endolongispinus* in comparison to other *Sodalis*- and *Symbiopectobacterium*-related endosymbionts. Endosymbionts are arranged in order of decreasing genome size as a proxy for the age of the symbiosis, as ordered as newest to oldest. ^1^Yuan et al., 2014; ^2^Martinson et al., 2020; ^3^Clayton et al., 2012; ^4^Oakeson et al., 2014; ^5^Toh et al., 2006; ^6^GenBank accession: GCA_001879235.1; ^7^Meseguer et al., 2017; ^8^Koga and Moran, 2014; ^9^GenBank accession: GCF_001602625.1; ^10^GenBank accession: GCF_900161835.1; ^11^GenBank accession: GCA_003668825.1; ^12^Husník and McCutcheon, 2016; ^12^Szabó et al., 2017.

| Genome | Status | Closest sequenced relative | Genome size | Coding density | Total predicted ORFs | Candidate pseudogenes/intact genes |
| --- | --- | --- | --- | --- | --- | --- |
| *Pectobacterium wasabiae*<sup>1</sup> | Opportunistic pathogen | *Pectobacterium atrosepticum* | 5.04 Mbp | 81% | 4,627 | 332/4316 |
| *Ca. Sym. endolongispinus* | Endosymbiont | *Pectobacterium wasabiae* | 4.65 Mbp | 51% | 5,864 | 2885/2993 |
| *Ca. Sym. sp. SyHa*<sup>2</sup> | Endosymbiont | *Pectobacterium wasabiae* | 4.56 Mbp | 57% | 5,510 | 2313/3237 |
| *Ca. Sym. sp. SyDa*<sup>2</sup> | Endosymbiont | *Pectobacterium wasabiae* | 3.64 Mbp | 72% | 3,795 | 842/3022 |
| *Ca. Sym. sp. SyCt*<sup>2</sup> | Endosymbiont | *Pectobacterium wasabiae* | 1.50 Mbp | 54% | 1,739 | 840/1013 |
| *Ca. Sym. sp. SyCl*<sup>2</sup> | Endosymbiont | *Pectobacterium wasabiae* | 246 Kbp | 64% | 289 | 107/197 |
| *Ca. Sodalis praecaptivus* HS<sup>3</sup> | Opportunistic pathogen | *Sodalis* sp. 159R | 5.16 Mbp | 77% | 4505 | 314/4204 |
| *Ca. Sodalis pierantonius* str. SOPE<sup>1,4</sup> | Endosymbiont | *Sodalis* HS | 4.51 Mbp | 46% | 5,288 | 3296/2418 |
| *Ca. Sodalis glossinidius*<sup>5</sup> | Endosymbiont | *Sodalis* HS | 4.31 Mbp | 52% | 5,750 | 3091/3025 |
| *Ca. Sodalis endolongispinus* | Endosymbiont | *Sodalis* HS | 3.59 Mbp | 53% | 4,368 | 1959/2453 |
| *Ca. Sodalis* sp. TME1<sup>6</sup> | Endosymbiont | *Sodalis* HS | 3.42 Mbp | 53% | 3,803 | 1596/2423 |
| *Ca. Sodalis* sp. SCIS<sup>9</sup> | Endosymbiont | *Sodalis* HS | 3.08 Mbp | 68% | 2,041 | 382/1681 |
| *Ca. Sodalis* sp. PSPU<sup>8</sup> | Endosymbiont | *Sodalis* HS | 2.23 Mbp | 68% | 2,041 | 382/1681 |
| *Ca. Sodalis* sp. PFLU<sup>9</sup> | Endosymbiont | *Sodalis* HS | 2.17 Mbp | 37% | 2,582 | 1896/1249 |
| *Ca. Sodalis* sp. HHAL<sup>10</sup> | Endosymbiont | *Sodalis* HS | 1.62 Mbp | 37% | 795 | 92/750 |
| *Ca. Sodalis* sp. BTRI<sup>11</sup> | Endosymbiont | *Sodalis* HS | 1.58 Mbp | 38% | 2,462 | 1285/1218 |
| *Ca. Gullenella endobia*<sup>12</sup> | Endosymbiont | *Sodalis* HS | 938 Kbp | 48% | 479 | 45/456 |
| *Ca. Doolittlea endobia*<sup>12</sup> | Endosymbiont | *Sodalis* HS | 847 Kbp | 58% | 658 | 160/570 |
| *Ca. Hoaglandella endobia*<sup>12</sup> | Endosymbiont | *Sodalis* HS | 637 Kbp | 78% | 530 | 33/510 |
| *Ca. Moranella endobia*<sup>12</sup> | Endosymbiont | *Sodalis* HS | 538 Kbp | 76% | 438 | 43/410 |
| *Ca. Mikella endobia*<sup>12</sup> | Endosymbiont | *Sodalis* HS | 353 Kbp | 74% | 282 | 26/267 |
| *Ca. Trabutinella endobia*<sup>13</sup> | Endosymbiont | *Sodalis* HS | 298 Kbp | 77% | 247 | 31/232 |

### Transposases recently proliferated within the genome of Sod. endolongispinus

*Sym. endolongispinus* and *Sod. endolongispinus* were screened for insertion sequences (ISs), which are types of mobile genetic elements in bacteria. ISs are typically made up of transposase genes along with other accessory and passenger genes (Mahillon and Chandler, 1998), and have previously been suggested to proliferate during the early stages of host restriction in endosymbionts (Plague *et al*., 2008; Gil *et al*., 2008; Belda *et al*., 2010; Schmitz-Esser *et al*., 2011; Clayton *et al*., 2012; Oakeson *et al*., 2014). *Sod. endolongispinus* encodes at least 220 transposase genes, 96% of which are part of the IS3 family (**Supplemental Figure 2A**). The rest of the transposases are part of the IS-NCY transposase family. Both of these IS families are encoded by the close relative *Sodalis* HS, but in smaller numbers and different proportions. The expansion of IS3 family transposases in *Sod. endolongispinus* appears to have occurred very recently, because the vast majority of these transposases are part of two distinct clusters of paralogs, where each cluster contains about 80 nearly identical copies of the same transposase that has proliferated throughout the genome (**Supplemental File 1**, **Supplemental Figure 3**). Only a handful of transposase duplications have a *dS* value (proxy for evolutionary divergence) greater than 0.5, which is the average dS of homologs between *Sod. endolongispinus* and *Sodalis* HS, suggesting that most transposition events occurred after divergence of the two species. In contrast, non-transposase gene duplicates, which comprise 101 genes, have an average *dS* of 1.3, suggesting that they likely duplicated prior to host restriction. Gene duplication prior to divergence is also supported by the fact that orthologs to most non-transposase duplicated genes are also encoded as duplicates on the genome of *Sodalis* HS. Long-term maintenance of gene duplicates is considered rare in prokaryotic genomes (Hooper and Berg, 2003); however, in certain bacterial species, gene duplicates do persist, and can accumulate to a considerable fraction of the genome (Gevers *et al*., 2004; Miller *et al*., 2011). In contrast to *Sod. endolongispinus*, we find that *Sym. endolongispinus* does not appear to have undergone an expansion of transposases.

Many of the identified transposase genes appear to have been pseudogenized in some way. In *Sod. endolongispinus* and *Sym. endolongispinus*, 26% and 70%, respectively, of all identified transposases have been flagged as pseudogenes nu the Pseudofinder software (**Supplemental Figure 3**). The vast majority of these pseudogene predictions are based on the shorter length of each transposase relative to the closest homologs available in NCBI. There are also some transposases that appear to have acquired nonsense mutations, and exist as multiple fragments on the genome. The fact that many transposases have become pseudogenized is not unique to the *P. longispinus* endosymbionts. Other *Sodalis* and *Symbiopectobacterium*-related symbionts show similar levels of pseudogenization among their transposases.

### Both gammaproteobacterial endosymbionts show complementary patterns of gene pseudogenization and loss in amino acid and vitamin biosynthesis

While the genomes of the gammaproteobacterial symbionts of *P. longispinus* are still large, the pseudogenization of nearly half of their genes allows us to ask whether gene inactivation events show nascent signals of the interdependency that is common in more established endosymbionts (Martin and Herrmann, 1998; Shigenobu, 200l; Wu *et al*., 2006; Gosalbes *et al*., 2008; McCutcheon and Moran 2010; Lamelas *et al*., 2011; Sloan and Moran, 2012; Husník *et al*., 2013; López-Madrigal *et al*., 2013; Bennett *et al*., 2014; Santos-Garcia *et al*., 2014; Luan *et al*., 2015; Husník and McCutcheon, 2016; Szabó *et al*., 2017; Ankrah *et al*., 2020). Clear patterns of complementary gene loss and retention have been observed in other mealybug symbioses that host intra-*Tremblaya* gammaproteobacterial symbionts, but in these other cases, the gammaproteobacterial endosymbionts have highly reduced and gene-dense genomes of less than 1 Mb, consistent with much longer periods of host restriction (Szabó *et al*., 2016; Husník and McCutcheon, 2016).

We find that *Sym. endolongispinus* and *Sod. endolongispinus* show signs of nascent complementarity in gene loss and retention. This pattern is most clear in key host-required pathways used to build essential amino acids and vitamins (**Figure 4A**). For example, the pathways for biosynthesis of the amino acids histidine, cysteine, arginine, threonine, methionine, and others, show signs of partitioning between both gammaproteobacterial genomes through reciprocal pseudogene formation and gene loss. Some of the biosynthetic genes missing or pseudogenized from both *Sod. endolongispinus* and *Sym. endolongispinus* (e.g. *ribC, bioA*, and *bioB*) are encoded either on the *Tremblaya* or on the host genome as bacterial HGTs (Husník and McCutcheon, 2016). There are also many pathway components that remain redundant in the system, with multiple gene copies present between the symbiotic partners (**Figure 4B**). For example, three of the genes responsible for lysine biosynthesis are encoded on the host as HGTs, but these genes are also retained in both *Sod. endolongispinus* and *Sym. endolongispinus*. There are also two instances where a required gene (*argA* [arginine] and *bioC* [biotin]) is missing completely from the symbiosis. These genes are also missing in older symbioses, and it is possible that their roles have been taken over by host proteins of eukaryotic origin (Husník *et al*., 2013).

**Figure 4:**
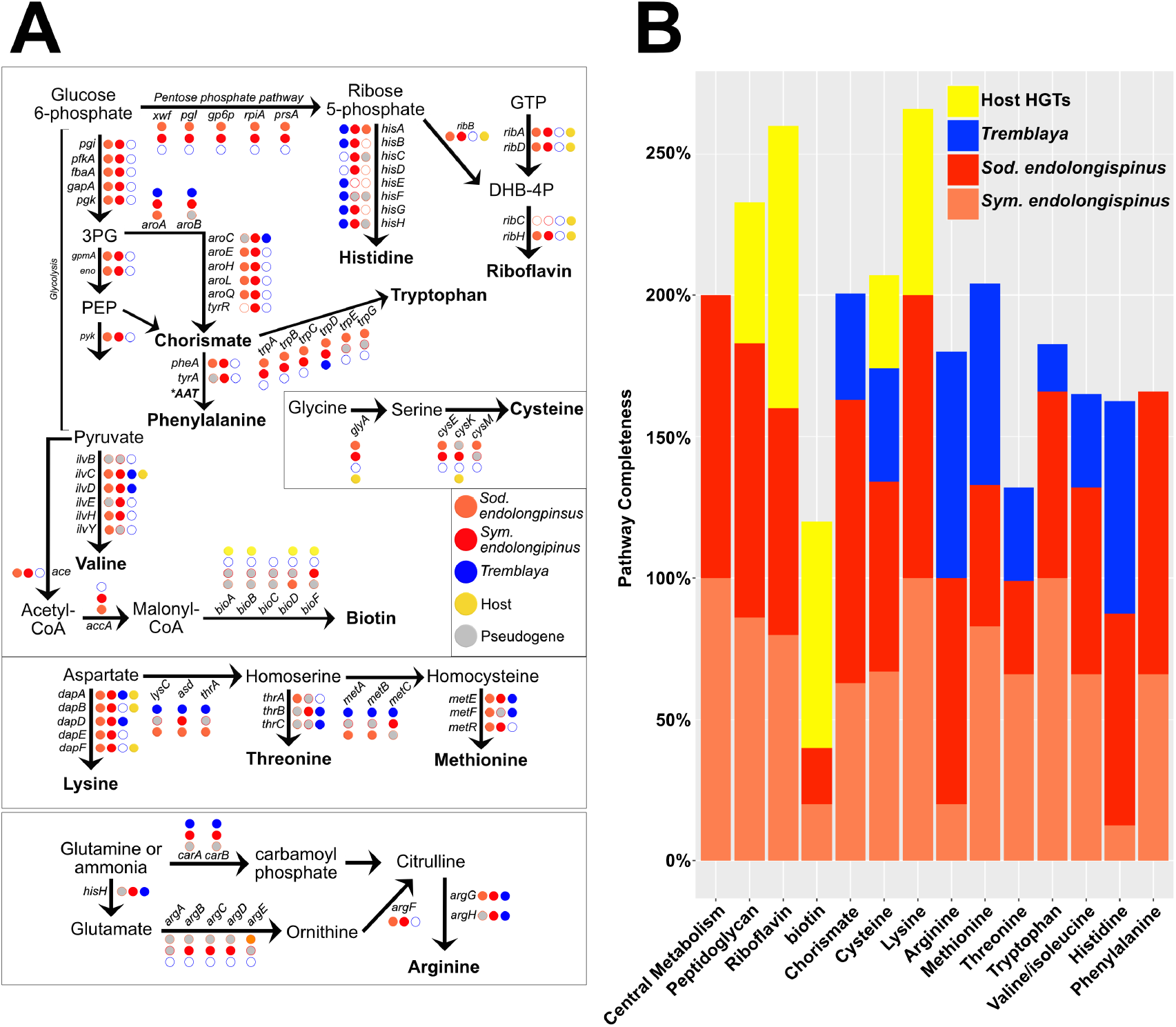
Distribution of metabolic genes in the P. longispinus symbiosis. **A)** Presence, absence, and pseudogenes among the various biosynthetic pathways in P. longispinus. Also shown are the central metabolism pathways (glycolysis, pentose phosphate, and acetate node). Pseudogenes are colored gray. The presence of a gene on the host genome (either native or from HGT) is shown as a filled yellow circle. **B)** Barplot showing percent completion of various metabolic pathways among the P. longispinus endosymbionts and host-encoded genes.

### Core metabolic and cell structural genes in gammaproteobacterial genomes are strongly retained

Contrary to the pattern of complementary degradation in pathways for amino acid and vitamin biosynthesis, genes that are part of the core metabolic and cell structural pathways show strong retention in both *Sod. endolongispinus* and *Sym. endolongispinus* (**Figure 4**). Specifically, genes for glycolysis, pentose phosphate, and the acetate node, as well as the other essential pathways (e.g. iron-sulfur cluster biosynthesis, tRNA modification), are completely intact on both of the young endosymbiont genomes in *P. longispinus*.

Finally, we investigated the pathway for peptidoglycan (PG) biosynthesis. PG is an important component of the bacterial cell envelope; it provides rigidity and shape to most bacterial cells (Otten *et al*., 2018). We have previously shown that in a related mealybug species, *Planococcus citri*, PG is produced by a biosynthetic pathway split between horizontally acquired genes encoded on the host genome and genes on the gammaproteobacterial endosymbiont genome (Bublitz *et al*., 2019). However, in *P. citri*, *Tremblaya* harbors an ancient and long-established gammaproteobacterial symbiont, *Ca*. Moranella endobia (hereafter, *Moranella*), which has a highly reduced genome with many deleted PG-related genes. *P. citri* and *P. longispinus* are somewhat closely related mealybugs and share the same PG-related bacterial HGTs on their nuclear genomes (Husník and McCutcheon, 2016). Here, we find that the core PG biosynthesis pathway is intact in *Sym. endolongispinus* (**Figure 5A**). In *Sod. endolongispinus*, however, two PG-related genes, *murF* and *ddl*, have acquired nonsense substitutions which fragment each gene into two separate ORFs (**Supplemental File 3**). Interestingly, these two genes are present as HGTs on the *P. longispinus* genome (Husník and McCutcheon, 2016), but it is unclear if these HGTs somehow complement the early loss of PG genes in *Sod. endolongispinus* in a manner similar to that in *P. citri* (Bublitz, *et al*., 2019).

**Figure 5:**
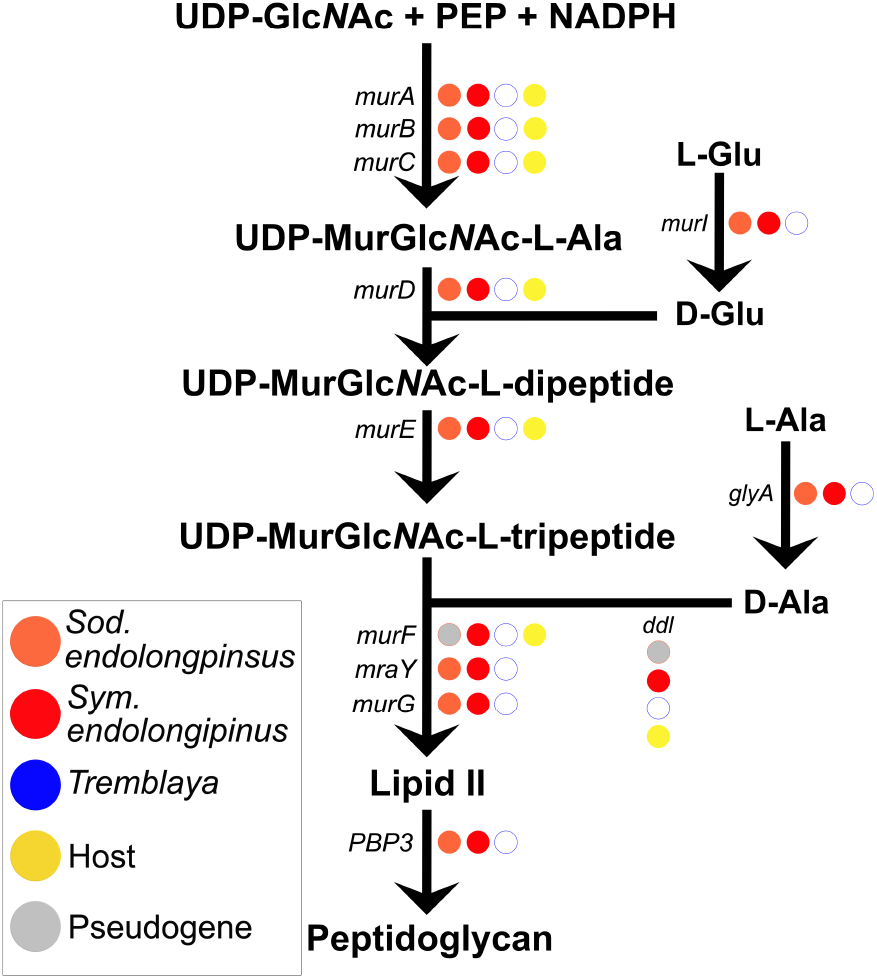
The peptidoglycan biosynthesis pathway in P. longispinus endosymbionts. Presence, absence, and pseudogenes within the peptidoglycan biosynthesis pathway in P. longispinus endosymbionts. Pseudogenes are colored gray. The presence of a gene on the host genome is shown as a filled yellow circle.

## Discussion

### Gammaproteobacterial endosymbionts in P. longispinus are of recent origin

We conclude that the gammaproteobacterial endosymbionts in *P. longispinus* mealybugs have been introduced into a host-restricted lifestyle relatively recently, on a timescale roughly similar to other young endosymbionts in insects and nematodes (Toh *et al*., 2006; Burke and Moran, 2011; Clayton *et al*., 2012; Oakeson *et al*., 2016; Boyd *et al*., 2016; Martinson *et al*., 2020). We base this conclusion on three features of their genomes. First, their genome sizes are large, comparable to those of free-living bacteria (**Table 1**, **Figure 3**) (Husník and McCutcheon, 2016), showing that they have not yet undergone most of the genome reduction seen in more established bacterial endosymbionts (McCutcheon and Moran, 2012). Second, they fall on relatively short branch lengths on phylogenomic trees relative to their non-endosymbiont relatives (**Figure 2**), indicating that they have not yet experienced the rapid sequence evolution typical of older endosymbiotic bacteria (Moran, 1996). Third, their GC contents at 4-fold degenerate sites in coding regions remains relatively high (**Supplemental File 2**), whereas older endosymbionts typically show pronounced AT biases at these sites (Wernegreen, 2002; Van Leuven and McCutcheon, 2012).

We attempted to infer which gammaproteobacterial endosymbiont might have been established first within *P. longispinus* bacteriocytes. The lower average *dS* and shorter branch length relative to its closest non-endosymbiotic relative (**Figure 2**) suggests that *Sod. endolongispinus* is the younger of the two gammaproteobacterial endosymbionts in *P. longispinus*. This is consistent with our rough estimate of 68,000 years as the divergence time between *Sod. endolongispinus* and *Sodalis* HS, compared with >100,000 years for the symbiosis events in the *Symbiopectobacterium* clade, as estimated by Martinson *et al*. (2020). It is important to emphasize that these dates are extremely speculative. Additionally, it is possible that the longer branch length of *Sym. endolongispinus* (as well as the other *Symbiopectobacterium* symbionts) relative to the non-endosymbiotic *Pectobacterium*/*Brenneria* spp. is due to the fact that a close relative to *Symbiopectobacterium* symbionts has not yet been sequenced.

### Creation of pseudogenes is likely coupled with rapid deletion

During their brief period of host restriction and vertical transmission, *Sym. endolongispinus* and *Sod. endolongispinus* have accumulated thousands of pseudogenes (**Figure 3**). This is in stark contrast to non-endosymbiotic bacteria, where pseudogenes have been reported to account for only 1-5% (in some cases, as high as 8%) of the genetic repertoire (Liu *et al*., 2004; Lerat and Ochman, 2005). The high level of gene inactivation in both *Sym. endolongispinus* and *Sod. endolongispinus* genomes is caused by many frameshifts and stop mutations, resulting in either truncated, run-on, and fragmented ORFs (**Supplemental Files 3-4**).

A small proportion of apparently functional genes appear to be cryptic pseudogenes, or genes that have elevated *dN/dS* values (> 0.3) relative to orthologs in a closely related non-endosymbiont genome (Clayton *et al*., 2012; Oakeson *et al*., 2014; Van Leuven *et al*., 2014; Burke and Moran, 2011). However, the vast majority of both intact and broken genes have low *dN/dS* values consistent with strong purifying selection. This suggests that pseudogenes in *Sym. endolongispinus* and *Sod. endolongispinus* have formed very recently and have not yet had time to accumulate substitutions that would elevate their *dN/dS* values. As noted in previous studies of pseudogene flux in several strains of *Salmonella* (Kuo and Ochman, 2010), and consistent with the previously reported deletional bias in bacterial genomes (Mira *et al*., 2001; Kuo and Ochman, 2009; Burke and Moran, 2011), it is likely that deletion of pseudogenes happens quickly, on time scales shorter than these genes can accumulate significant numbers of sequence substitutions.

### Recent transposase expansion a common, but not universal, feature of early genomic disruption in endosymbionts

*Sod. endolongispinus* encodes 220 transposases, over an order of magnitude greater than its closest free-living relative. The high number of transposases in *Sod. endolongispinus* is consistent with previous reports of IS expansion as a byproduct of relaxed selection on large parts of the genome, as well as a mechanism for genome rearrangement and reduction (Plague *et al*., 2008; Belda *et al*., 2010; Schmitz-Esser *et al*., 2011; Oakeson *et al*., 2014; Mahillon Chandler, 1998; Hendry *et al*., 2018). Indeed, other recently acquired *Sodalis* endosymbionts also encode high numbers of transposases (Clayton *et al*., 2012; Oakeson *et al*., 2016). Each *Sodalis* endosymbiont encodes different types and distributions of IS families (**Supplemental Figure 2A**). Certain transposase families present in the other *Sodalis* endosymbionts encode transposases that are not found in *Sodalis* HS (e.g. IS21 in SOPE, IS1 in *Sodalis* sp. SCIS). Given that ISs are highly dynamic, moving within and between genomes (Touchon *et al*., 2007), it is possible that even a very close relative to *Sodalis* HS could have radically different types and distributions of ISs.

Similar to *Sod. endolongispinus*, SOPE and *S. glossinidius* have likely undergone an IS expansion very recently, since the vast majority of the transposase sequences fall into only a handful of clusters of paralogs composed of nearly identical copies of just a few families per genome (**Supplemental File 1**, **Supplemental Figure 2B**), consistent with recent analyses of SOPE (Gil *et al*., 2008; Oakeson *et al*., 2014). The dearth of genetic diversity among transposases in each genome suggests that, like pseudogenes, transposases are rapidly deleted from the genome after a burst of transposition at the onset of endosymbiosis. Because transposases are known for facilitating deletions (Mahillon and Chandler, 1998) and proposed to be involved in genome reduction in endosymbionts (Siguier *et al*., 2014), it is possible that this property of transposases ultimately results in the elimination of IS elements themselves (Plague *et al*., 2008; Schmitz-Esser *et al*., 2011; Siguier *et al*., 2014). Consistent with this idea, the genomes of older endosymbionts encode low numbers, if any, IS elements (**Supplemental Figure 2**).

IS proliferation does not appear to be a universal phenomenon in young endosymbionts, at least not among the *Sodalis* and *Symbiopectobacterium* endosymbionts screened here. Several *Sodalis* endosymbionts (e.g. *Sodalis* sp. TME1, *Sodalis* sp. SCIS), whose genomes are comparable in size to *Sod. endolongispinus* (**Table 2**), appear to have few or no transposases (**Supplemental Figure 2**). None of the *Symbiopectobacterium* symbionts encode nearly as many transposases as *Sod. endolongispinus* or *S. glossinidius*, but have similar or lower numbers of transposases relative to *P. wasabiae* and *P. cartovorum*, the closest non-endosymbiotic relatives of the *Symbiopectobacterium* clade. The relative lack of transposases is surprising, given the vast amount of superfluous genome space in these young endosymbiont genomes (*Table 2*), which should support IS proliferation similar to what is observed in *Sod. endolongispinus* and SOPE. It seems likely that there are large and diverse populations of free-living *Sodalis* and *Symbiopectobacterium* strains in nature that vary in IS content, and that the amount of IS proliferation that occurs in a newly established endosymbiont reflects the IS load of the particular strain of free-living bacterium from which it was derived.

It is also possible that the dearth of detectable transposase genes is caused by the low assembly quality of some of the endosymbiont genomes that are highly fragmented (e.g. >100 contigs) (**Supplemental Table 1**). The highly fragmented nature of these short-read assemblies can, in part, be caused by the prevalence of identical or nearly identical transposases that cannot be resolved using short reads alone. For example, the assembly corresponding to *Sym. endolongispinus* published by Martinson *et al*. (2020) is comprised of 83 contigs (using only the Illumina reads published by Husník and McCutcheon [2016]), with only 6 transposases detected using the ISfinder software that’s included in the Prokka annotation pipeline. Our hybrid assembly using PacBio and Illumina reads resolved this genome into 9 contigs, from which we were able to identify 36 transposases. Similarly, our Illumina-only assembly (using Unicycler) of the *Sod. endolongispinus* genome resulted in 109 contigs, with only 7 identifiable transposases; our PacBio/Illumina hybrid assembly resolved the *Sod. endolongispinus* genome into 3 contigs, with 220 identifiable transposases. It seems likely that the estimated number of transposases in genomes generated from short-read data alone are significantly underestimated, with near-identical ISs collapsing into small unassembled contigs.

### Establishment of interdependence occurs early during endosymbiont establishment

Previous research has demonstrated that newly evolved endosymbiotic bacteria lose genes in response to the pre-existing genetic inventory of their co-symbionts and (if present) HGTs on the host genome (Wu *et al*., 2006; McCutcheon and Moran, 2007; Nikoh and Nakabachi, 2009; McCutcheon *et al*., 2009; Sloan and Moran, 2012; Husník *et al*., 2013; Luan *et al*., 2015; Nowack *et al*., 2016). The occurrence of two recently acquired endosymbionts in *P. longispinus* presented us with a unique opportunity to investigate the inception of genomic complementarity and metabolic interdependence in a complex four-way symbiosis.

During this early period of host restriction, we might expect to see more rapid gene loss in pathways whose precursors, intermediates, and products are more easily transported between different members of the symbiosis. This can include metabolites for which dedicated transporters (e.g. amino acid permeases) already exist in the free-living predecessor. For example, *Sod. endolongispinus* encodes genes for the transport of histidine and biotin, possibly contributing to the rapid loss of the biotin and histidine biosynthesis pathways in that endosymbiont (**Figure 4**). Many genes that are part of amino acid and vitamin metabolism are either encoded on the host’s nuclear genome as HGTs from bacteria, or on *Tremblaya*’s diminutive genome, relieving the need for these new gammaproteobacterial symbionts to continue maintaining these genes. Consistent with this, we observe loss and pseudogenization of many pathway components for amino acid and vitamin biosynthesis in *Sym. endolongispinus* and *Sod. endolongispinus* (**Figure 4A**), suggesting that genes in these pathways are lost rapidly and in response to genes already present in the symbiosis.

As *Sym. endolongispinus* and *Sod. endolongispinus* are relatively new to a host-dependent lifestyle, they still encode many genes redundant with other genes present in the system (**Figure 4B**). Most of these redundant genes appear to be undergoing strong purifying selection, evident from their low *dN/dS* values (**Supplemental Files 3-4**). Because other older and longer-established mealybug symbioses show little evidence of genetic redundancy across genomes (Husník and McCutcheon, 2016), we suspect that many of the redundant genes in the gammaproteobacterial endosymbionts simply have not had a chance to accumulate substitutions that would break genes, elevate their *dN/dS* values, or delete them completely from one genome or the other.

### Slower loss of core metabolic and structural genes

In contrast to the rapid gene loss in pathways for amino acid and vitamin biosynthesis, genes in pathways for peptidoglycan biosynthesis and central metabolism are more strongly conserved in both of the gammaproteobacterial symbionts of *P. longispinus*. We have previously demonstrated that in *Moranella*, the ancient gammaproteobacterial symbiont of *Planococcus citri*, peptidoglycan biosynthesis occurs in concert with bacterial genes encoded on its host’s nuclear genome as HGTs (Bublitz *et al*., 2020). Consequently, *Moranella* has lost much of its peptidoglycan biosynthesis pathway and is presumably reliant on the import of host-derived proteins. While this level of cellular integration represents a potential future state for *Sym. endolongispinus* and *Sod. endolongispinus*, it appears that this level of integration has not yet been achieved in the relatively short amount of time that these symbionts have been inhabiting *P. longispinus*.

We hypothesize that the generally stronger retention of the peptidoglycan biosynthesis pathway throughout the early stages of endosymbiosis is due to the difficulty of importing host-derived proteins, such as MurF (Bublitz *et al*., 2020), into the newly established gammaproteobacterial endosymbionts. The loss of two key components of the peptidoglycan biosynthesis pathway (*murF* and *ddl*) in *Sod. endolongispinus* is, therefore, surprising (**Figure 5**). It is possible that the recurrent recruitment and maintenance of endosymbionts from the *Sodalis* genus (Husník and McCutcheon, 2016) has made mealybugs particularly suited for rapid cellular integration of *Sodalis* relatives after infection. This is consistent with stronger conservation of peptidoglycan biosynthesis in *Sym. endolongispinus*, even though we estimate that *Sym. endolongispinus* is the older of the two gammaproteobacterial endosymbionts. Members of the *Symbiopectobacterium* clade do not seem to commonly infect mealybugs, as only one of seven mealybug species for which we have genomic data house a *Symbiopectobacterium*-related symbiont (Husník and McCutcheon, 2016; Szabó *et al*., 2016). It also possible that, since the loss of unnecessary or redundant genes is a stochastic process, and given that both gammaproteobacterial symbionts are relatively young, the observed pattern of loss and retention in *Sod. endolongispinus* and *Sym. endolongispinus* is simply due to chance.

## Acknowledgements

We thank Diane Brooks, Paul Caccamo, Filip Husník, Genevieve Krause, Piotr Łukasik, Mitch Syberg-Olsen, Dan Vanderpool, Catherine Armbruster, and Travis Wheeler for helpful discussions and insights during the course of this project. We thank the Caltech Kavli Nanoscience Institute for maintenance of the TF-30 electron microscope. This work was supported by the National Science Foundation (IOS-1553529; to JPM), the Gordon and Betty Moore Foundation (GBMF5602; to JPM), the National Aeronautics and Space Administration Astrobiology Institute (NNA15BB04A; to JPM), and the National Institute of Allergy and Infectious Diseases (2 P50 AI150464; to PJB).

---

**Supplemental Figure 1:**
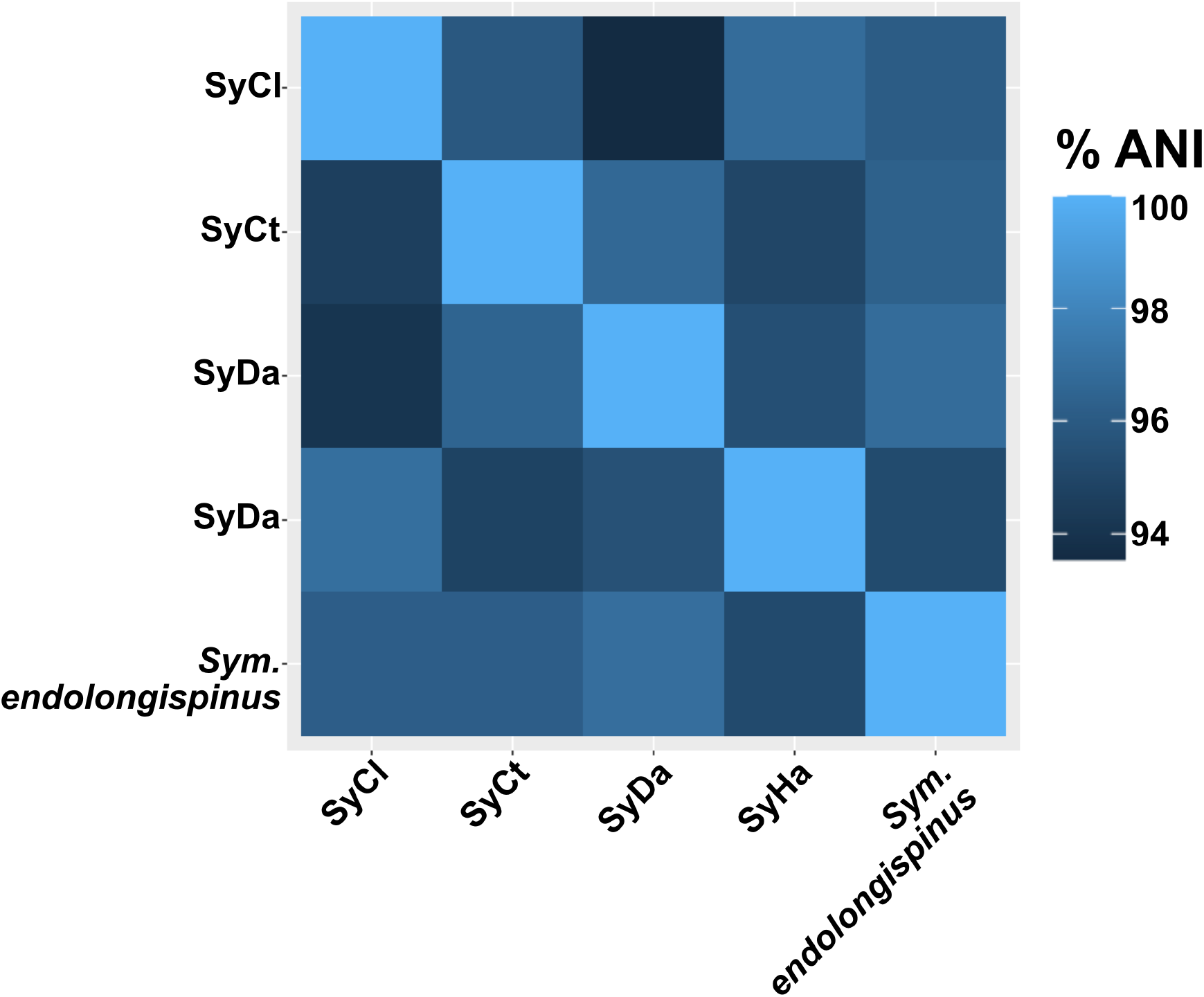
Pairwise average nucleotide identity, estimated using a BLAST-based best-hits approach

**Supplemental Figure 2:**
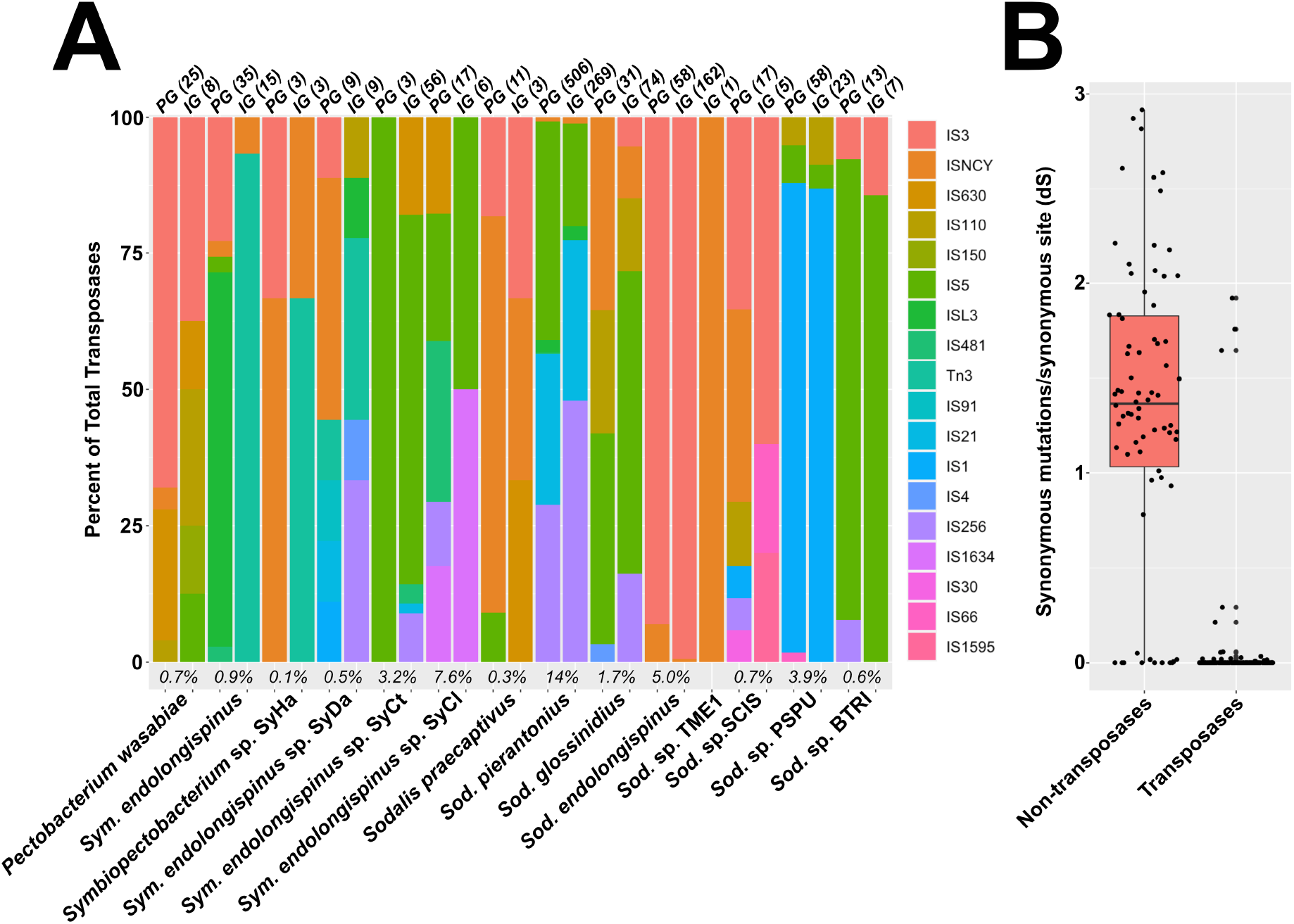
**A)** Numbers and distributions of various transposase families found in the endosymbiont genomes that are discussed in this study. The italicized numbers at the bottom of each stacked barplot denote the prevalence of transposase in each study, expressed as percent of total genes. The absolute numbers of intact (IG) and pseudogenized (PG) transposases in each genome is shown above each bar. Sodalis sp. TME1 only has a single transposase encoded on its genome (0.03% of total genes encoded by that endosymbiont) **B)** Boxplots demonstrating the generally low (mostly zero) dS values of duplicated transposases (n=151), compared with other gene duplicates (n=101) in Sod. endolongispinus.

**Supplemental Figure 3:**
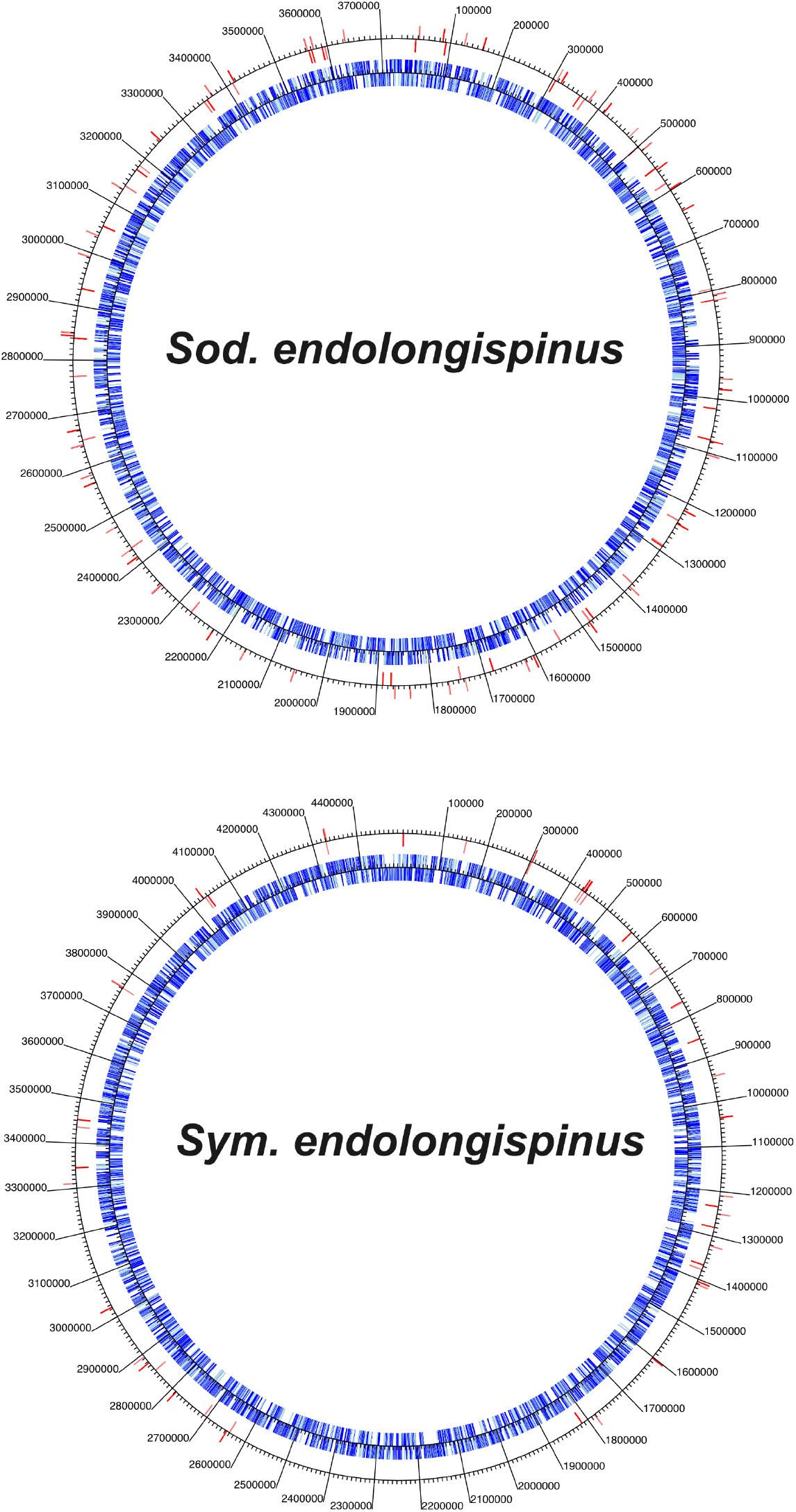
Location of transposases (colored red on the outside track) in the genomes of Sod. endolongispinus and Sym. endolongispinus. Input genes are shown blue on the inside track.

**Supplemental Table 1.**
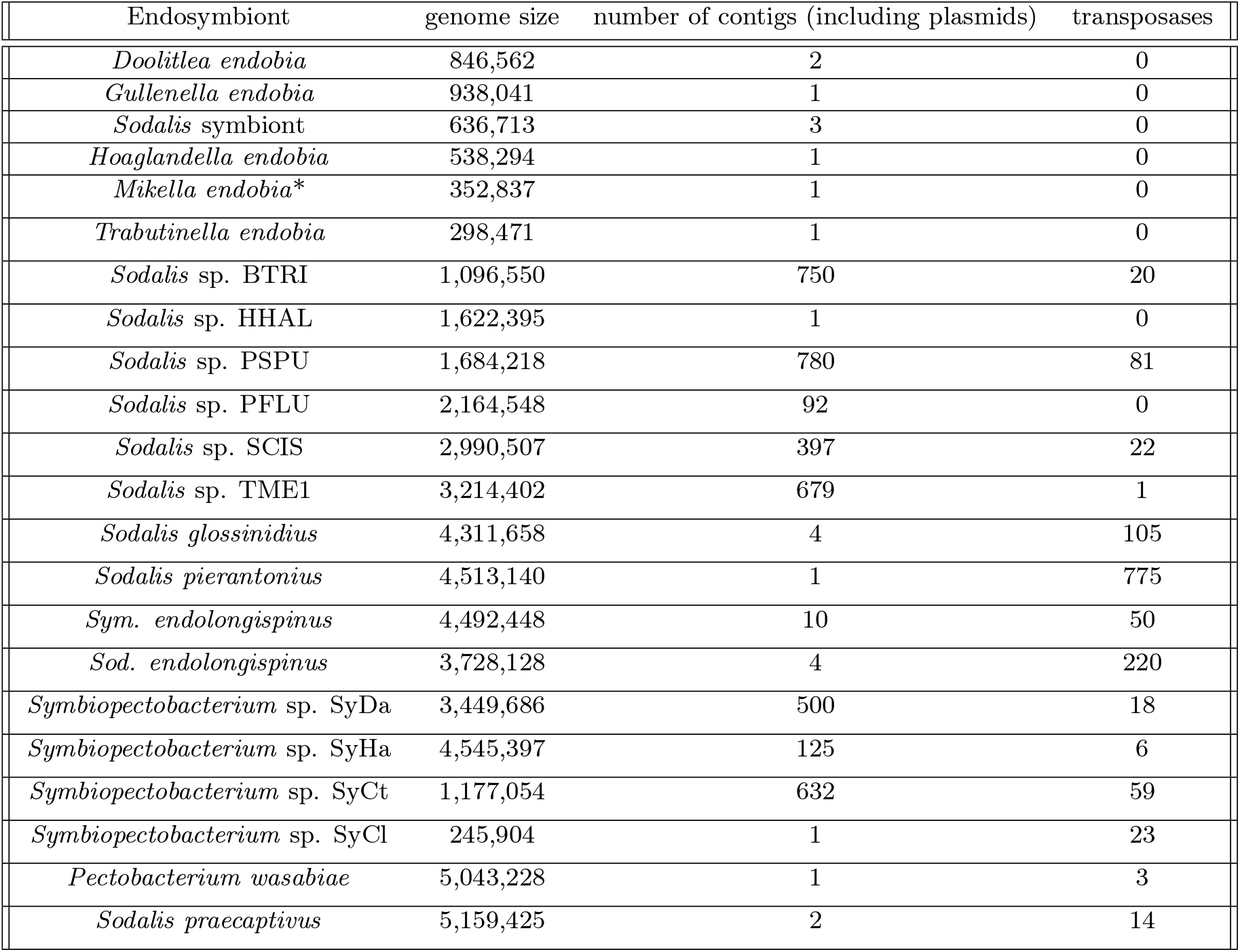
Assembly information of *Symbiopectobacterium*- and *Sodalis*-related endosymbiont genomes, in the context of the number of transposases identified in each genome.

| Endosymbiont | genome size | number of contigs (including plasmids) | transposases |
| --- | --- | --- | --- |
| *Doolitlea endobia* | 846,562 | 2 | 0 |
| *Gullenella endobia* | 938,041 | 1 | 0 |
| *Sodalis* symbiont | 636,713 | 3 | 0 |
| *Hoaglandella endobia* | 538,294 | 1 | 0 |
| *Mikella endobia* * | 352,837 | 1 | 0 |
| *Trabutinella endobia* | 298,471 | 1 | 0 |
| *Sodalis* sp. BTRI | 1,096,550 | 750 | 20 |
| *Sodalis* sp. HHAL | 1,622,395 | 1 | 0 |
| *Sodalis* sp. PSPU | 1,684,218 | 780 | 81 |
| *Sodalis* sp. PFLU | 2,164,548 | 92 | 0 |
| *Sodalis* sp. SCIS | 2,990,507 | 397 | 22 |
| *Sodalis* sp. TME1 | 3,214,402 | 679 | 1 |
| *Sodalis glossinidius* | 4,311,658 | 4 | 105 |
| *Sodalis pierantonius* | 4,513,140 | 1 | 775 |
| *Sym. endolongispinus* | 4,492,448 | 10 | 50 |
| *Sod. endolongispinus* | 3,728,128 | 4 | 220 |
| *Symbiopectobacterium* sp. SyDa | 3,449,686 | 500 | 18 |
| *Symbiopectobacterium* sp. SyHa | 4,545,397 | 125 | 6 |
| *Symbiopectobacterium* sp. SyCt | 1,177,054 | 632 | 59 |
| *Symbiopectobacterium* sp. SyCl | 245,904 | 1 | 23 |
| *Pectobacterium wasabiae* | 5,043,228 | 1 | 3 |
| *Sodalis praecaptivus* | 5,159,425 | 2 | 14 |

